# Growth Factors do not regulate Golgi Complex-to-ER relocation of GalNAc-Ts in HeLa cells

**DOI:** 10.1101/071225

**Authors:** Gaetan G. Herbomel, Raul E. Rojas, Duy T. Tran, Monica Ajinkya, Lauren Beck, Lawrence A. Tabak

## Abstract

Mucin-type O-glycosylation is initiated by the UDP-GalNAc polypeptide:*N*-acetylgalactosaminyltransferase (GalNAc-T) family of enzymes. Their activity results in the GalNAc α1-O-Thr/Ser structure, termed the Tn antigen, which is further decorated with additional sugars. In neoplastic cells, the Tn antigen is often overexpressed. Because O-glycosylation is controlled by the activity of GalNAc-Ts, their regulation is of great interest. Previous reports suggest that growth factors, EGF or PDGF, induce Golgi complex-to-endoplasmic reticulum (ER) relocation of both GalNAc-Ts and Tn antigen in HeLa cells, offering a mechanism for Tn antigen overexpression termed “GALA”. However, we were unable to reproduce these findings. Upon treatment of HeLa cells with either EGF or PDGF we observed no change in the co-localization of endogenous GalNAc-T1, GalNAc-T2 or Tn antigen with the Golgi complex marker TGN46. There was also no enhancement of localization with the ER marker calnexin. We conclude that growth factors do not cause redistribution of GalNAc-Ts from the Golgi complex to the ER in HeLa cells.

## Introduction

Mucin-type O-glycans decorate a wide range of secreted and transmembrane proteins. These molecules play a diverse set of physiological roles including structural roles within the extracellular matrix (Zhang et al., 2010), the function of the innate immune system (Tabak, 1995), and involvement in key signal transduction pathways (Kufe, 2013). In common with other secreted and trans-membrane proteins, the polypeptide core of mucins is synthesized in the endoplasmic reticulum (ER) and then transported to the Golgi complex via the ER-Golgi complex intermediate compartment (ERGIC). A family of enzymes, termed the UDP-GalNAc polypeptide:N acetylgalactosaminyltransferases (GalNAc-Ts, EC 2.4.1.41) initiates the addition of the first sugar in O-glycan biosynthesis in the Golgi complex by catalysing transfer of N-acetylgalactosamine (GalNAc) from the sugar donor UDP-GalNAc to the hydroxyl group of Thr/Ser residues of the protein core (Bennett et al., 2012). The resultant structure GalNAc α 1-O-Thr/Ser is termed “Tn antigen”. Normally, additional sugars are added stepwise, resulting in an extraordinary array of diverse O-glycan structures (Bennett et al., 2012). In contrast, Tn antigen is often expressed in neoplastic cells (Brockhausen, 2006; Itzkowitz et al., 1991; Konno et al., 2002; Konska et al., 2006; Laack et al., 2002; Springer, 1984), and is thought to contribute to their “invasiveness” (Radhakrishnan et al., 2014).

Mucin-type O-glycans are often arrayed in clusters due to variably glycosylated repeating Thr/Ser-containing sequences that form the sites of sugar attachment. Further, recent O-glycoproteomic studies (e.g. Steentoft et al., 2013) reveal that many non-mucinous secretory proteins are O-glycosylated, often with isolated glycosites, and these sites may have profound effect on protein functions. Thus, the acquisition, pattern, and density of O-glycans may be regulated by both the substrate specificity and the spatiotemporal expression of the multiple GalNAc-Ts. Detailed studies of GalNAc-T substrate specificity point to a requirement for either unique GalNAc-Ts, or multiple GalNAc-Ts working in concert, to ensure that the acquisition of O-glycans is complete (Bennett et al., 2012; Gerken et al., 2013; Revoredo et al., 2016; Steentoft et al., 2013).

Gill et al. (2010) reported that either EGF or PDGF activation of the Src-kinase pathway stimulates O-glycosylation initiation in HeLa cells. Their model posits that recruitment of active Src family kinases (SFKs) to the Golgi complex modulates the formation of transport carriers formed by the action of the GTPases, Arf1 and its effector the COPI complex, the molecular machinery known to control retrograde transport of Golgi complex resident proteins to the ER (Spang, 2013). Using *Helix pomatia* (HPA) lectin to detect Tn antigen, Gill et al. (2010) observed progressive re-distribution of HPA staining from the Golgi complex to the surrounding cytoplasm following growth factor treatment. To account for these findings, they suggested that GalNAc-Ts (but not other glycosyltransferases) located in the Golgi complex are uniquely sorted into retrograde carriers that transport these enzymes to ER resulting in higher levels of the Tn antigen in the ER of cells stimulated with growth factors compared to control cells. They term this novel mechanism for Tn antigen overexpression “GALA”.

This model has important implications for the regulation of mucin biosynthesis. If O-glycan acquisition were initiated in the ER due to selective re-direction of GalNAc-Ts, this could result in a more densely glycosylated Tn antigen since the remaining O-glycosylation machinery is not available until the mucin traffics to the Golgi complex. O-glycan density underlies many biological roles including enhancement of protein resistance to proteolysis (van der Post et al., 2013; Zhang et al., 2014), as well as imparting important structural properties of some surface receptors (Jentoft, 1990; Ostuni et al., 2014; Paszek et al., 2014), and the selectivity with which the mucin-type protein interacts with lectins (Dam et al., 2007; Kato et al., 2008). Moreover, in vitro studies suggest that the presence of Tn antigen inhibits extension of neighbouring glycans by the relevant glycosyltransferases (Brockhausen et al., 2009). Moreover, it is known that ER localized O-mannosylation influences the subsequent addition of GalNAc by GalNAc-Ts (Tran et al., 2012); if GalNAc-Ts re-localize to the ER, it is plausible that this could influence the acquisition of O-mannose residues. Therefore, if GalNAc-Ts re-locate to the ER, this would offer an important mechanism to alter the sugar coat of cell surfaces and their functions. Given the potential implications suggested by Gill et al. (2010) we attempted to reproduce several of the key observations published by Gill and colleagues (Gill et al., 2010) as a starting point to conducting studies that would build upon these findings. However, when we performed co-localization studies with experimental approaches similar to those reported by Gill et al., 2010, and additional ones using standard ER and Golgi complex markers in combination with antibodies to GalNAc-T1 and -T2, in EGF- or PDGF-stimulated HeLa cells (Gill et al., 2010; Gill et al., 2013), we failed to obtain any evidence for changes in the normal localization of endogenous GalNAc-T1 and GalNAc-T2. Rather, GalNAc-Ts remain predominately localized to the Golgi complex. Moreover, we did not detect changes in either the location of HPA-lectin reactive materials. Taken together, our experiments lead us to conclude that in HeLa cells, activation by the growth factors EGF and PDGF does not stimulate O-glycosylation initiation of mucins via the relocation of GalNAc-T from the Golgi complex to the ER.

## Results

### Growth factor stimulation has no effect on the Golgi complex localization of GalNAc-Tl and -T2

To determine if growth factor treatment effects the localization of endogenous GalNAc-T1 and -T2 we treated HeLa CCL-2 cells (acquired from ATCC) with EGF or PDGF, using conditions previously reported as sufficient to cause redistribution of GalNAc-T from the Golgi complex (Gill et al., 2010). GalNAc-T1 and -T2 are the most abundant GalNAc-T transcripts expressed in HeLa cells and thus represented a logical choice for these studies (Fig. S1). To directly visualize if GalNAc-Ts were located in the Golgi complex or ER, cells from all treatment conditions were stained with antibodies to TGN-46 (a Trans-Golgi complex marker), calnexin (an ER marker) and either endogenous GalNAc-T1 or endogenous GalNAc-T2 (the kind gift of Dr. Ulla Mandel, U. Copenhagen). In serum starved cells that were not treated with additional growth factors both GalNAc-T1 (Fig. 1A, A′-D′) and GalNAc-T2 (Fig. 1B, A′-D′) co-localized with TGN46. When serum starved cells were treated with EGF for 4 hours or PDGF for 3 hours both GalNAc-T1 (Fig. 1A, E′-L′) and GalNAc-T2 (Fig. 1B, E′-L′) remained co-localized with TGN46. To quantify the proportion of GalNAc-Ts located in the Golgi complex versus the ER under the various treatment conditions we determined the Manders’ correlation coefficient (MCC) (Dunn et al., 2011; Manders et al., 1993; McDonald and Dunn, 2013) of the GalNAc-T/TGN46 and GalNAc-T/calnexin staining, respectively (Fig. 1C). We found a strong coincidence of endogenous GalNAc-T1 or -T2 with TGN46 in both control, serum starved, cells and those stimulated with EGF or PDGF with a MCC of 0.9 ± 0.1. In contrast, the MCC between either GalNAc-T1 or -T2 and calnexin was 0.4 ± 0.1 under all conditions tested. Together, our data demonstrates that EGF and PDGF treatment of serum starved HeLa cells does not alter the Golgi complex localization of GalNAc-T1 or -T2.

**Figure 1.**
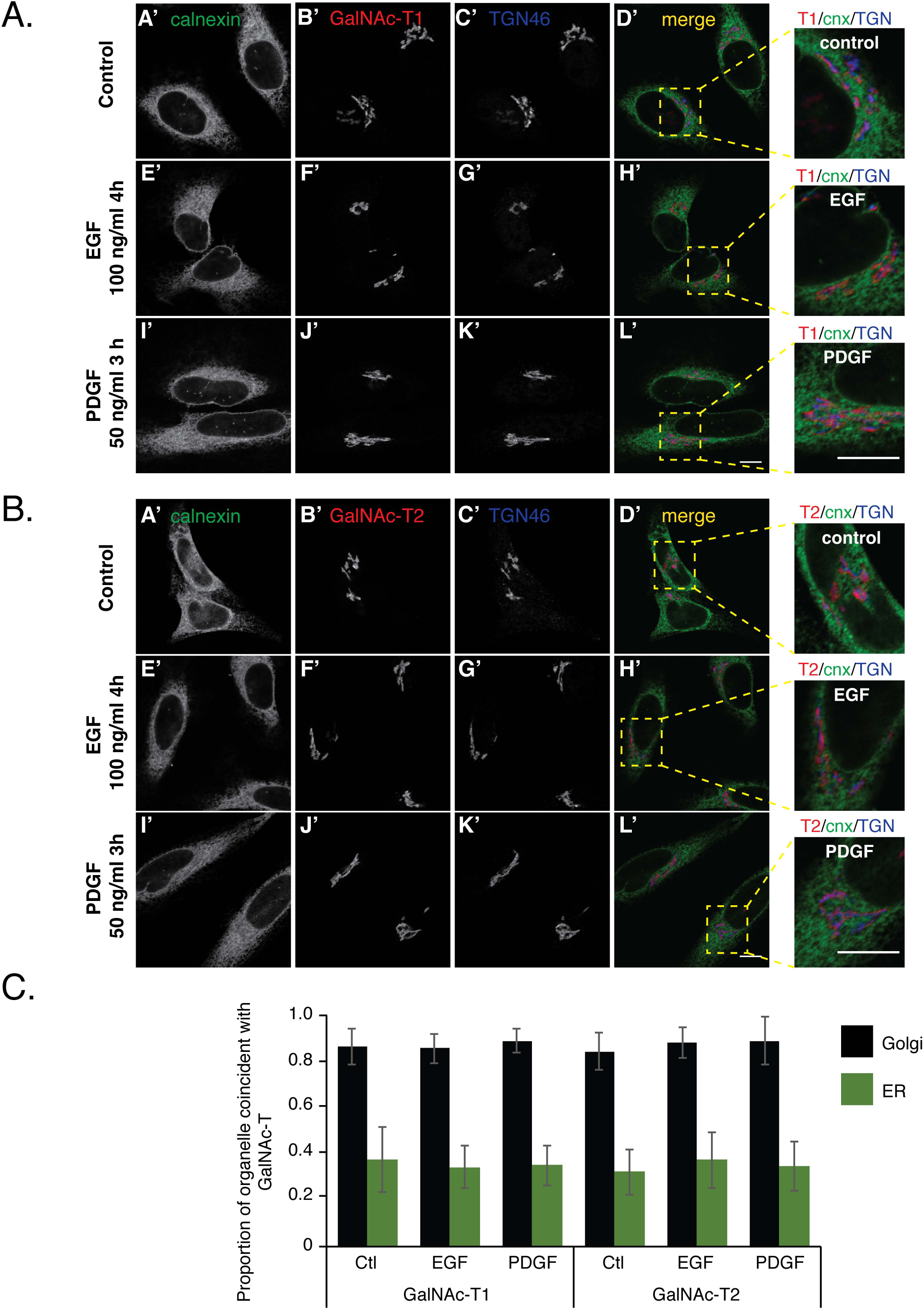
Growth factor treatment does not change Golgi complex localization of GalNAc-T1 or -T2. Serum starved HeLa cells were stimulated with 100 ng/ml of EGF for 4 h, 50 ng/ml of PDGF for 3 h or left untreated as a control. Cells were subsequently immunostained with antibodies to the Golgi complex marker TGN46 (AC′, AG′, AK′, BC′, BG′ and BK′), the ER marker calnexin (cnx) (AA′, AE′, AI′, BA′, BE′, and BI′) and either endogenous GalNAc-T1 (AB′, AF′, and AJ′) or GalNAc-T2 (BB′, BF′ and BJ′). Merged channels (AD′, AH′, AL′, BD′, BH′, and BL′) demonstrate that growth factor treatments have no effect on GalNAc-T1 or -T2 Golgi complex localization. Insets depicting a magnified view of merged imaging channels. Individual confocal sections shown in (A) are representative of 95, 74 and 78 cells for Control, EGF and PDGF treatments, respectively. Images in (B) are representative of 73, 79, and 61 cells for Control, EGF and PDGF treatments, respectively. Scale bars, 10 *μ*m. (C) Manders’ correlation coefficient was used to quantitate the degree of coincidence between GalNAc-T1 or -T2 with TGN46 (Golgi complex) and GalNAc-T1 or -T2 with calnexin (ER) in control, EGF and PDGF treated cells from confocal sections acquired over the entire volume of each cell. **Values represent the mean ± S.D**. from the number of cells described in (A) and (B).

### Proteins containing Tn antigen remain localized within the Golgi complex after growth factor stimulation

Because our growth factor treatment did not result in a change in GalNAc-T localization we next determined if EGF or PDGF treatment of serum starved HeLa cells resulted in a change in Tn antigen localization as was also reported by Gill et al. 2010. We similarly visualized the Tn antigen using a fluorescently labelled HPA (also referred to as HPL) lectin but additionally co-stained for both a Golgi complex (TGN46) and an ER (calnexin) marker. After treatment of serum starved cells with either EGF for 4 hours (Fig. 2A, E′-H′) or PDGF for 3 hours (Fig. 2A, I′-L′) HPA-reactive material was found localized with TGN46, similar to serum starved control cells (Fig. 2A, A′-D′). To quantitate the proportion of HPA-reactive material found in either the Golgi complex or ER we calculated the MCC of HPA-reactive material/TGN46 or HPA-reactive material/calnexin, respectively. We found a strong coincidence of HPA-reactive material with the Golgi complex with a MCC of 0.8 ± 0.1 for untreated and cells treated with EGF or PDGF, while the coincidence of HPA-reactive material with ER was low with a MCC of 0.3 ± 0.1, that did not change after growth factors treatment (Fig. 2B). Together, our data indicate that EGF or PDGF treatment of serum starved HeLa cells has no effect on the Golgi complex localization of proteins carrying the Tn antigen.

**Figure 2.**
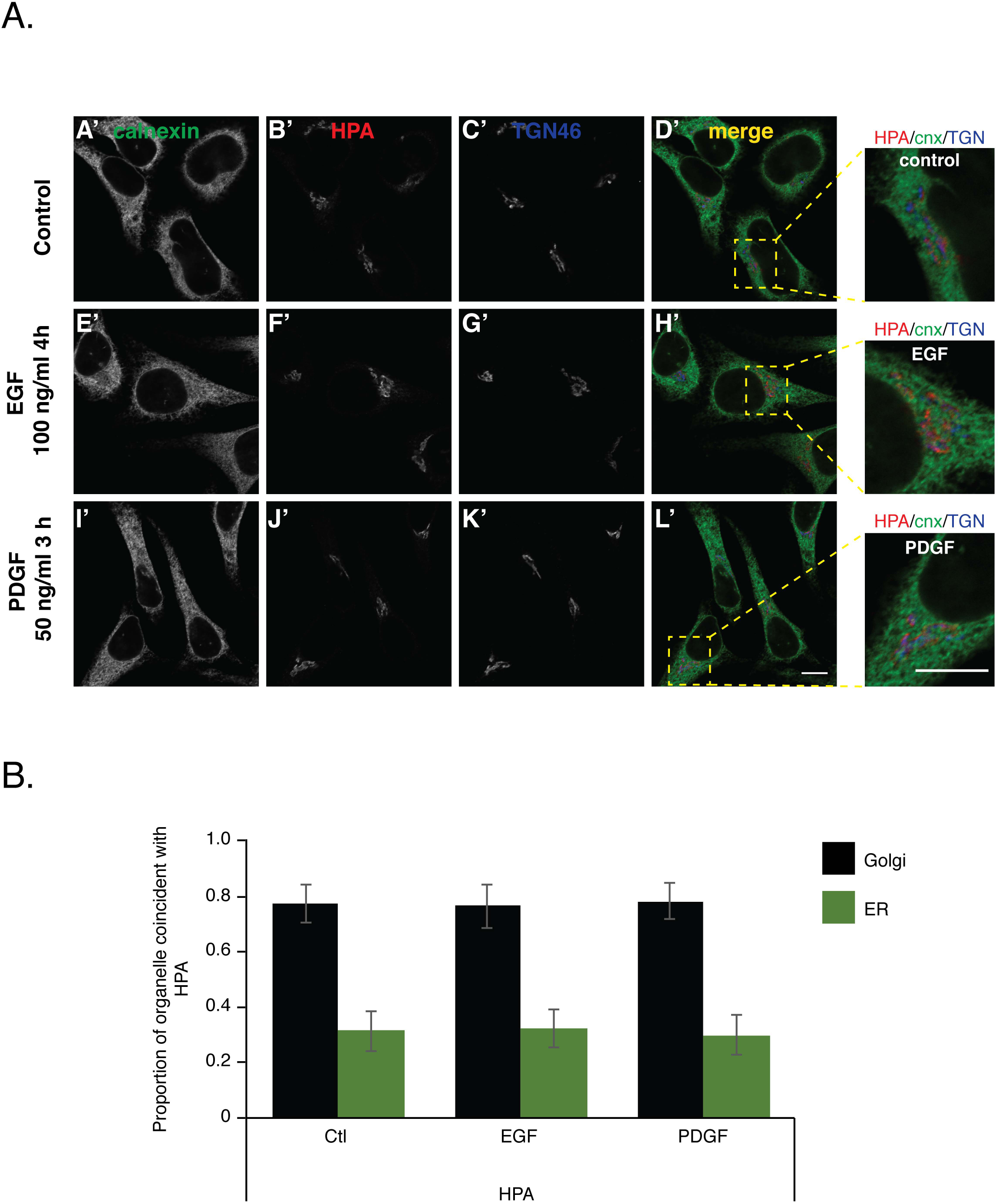
Growth factor treatment has no effect on Tn antigen localization. Serum starved HeLa cells were either left unstimulated, treated with 100 ng/ml of EGF for 4 h or 50 ng/ml of PDGF for 3 h. Cells were subsequently immunostained with antibodies to TGN46 (AC′, AG′ and AK′), calnexin (AA′, AE′, and AI′) and the lectin HPA (AB′, AF′ and AJ′). Merged channels (AD′, AH′, and AL′) show that neither EGF nor PDGF treatment move Tn antigen away from the Golgi complex. Individual confocal sections shown are representative of 93, 76 and 71 cells for Control, EGF and PDGF treatments, respectively. Scale bars, 10 *μ*m. (B) To quantitate the amount of Tn antigen found in Golgi complex versus ER we calculated the Manders’ correlation coefficient for HPA/TGN46 and HPA/calnexin signals acquired over the volume of individual cells, respectively. **Values represent the mean ± S.D**. of quantified experimental replicates described in (A).

### Growth factor treatment results in rapid signalling in HeLa cells

Our results indicated that EGF or PDGF treatment of serum starved cells does not result in any changes in GalNAc-T or Tn antigen localization away from the Golgi complex (Fig. 1 and 2). Because these results were contrary to those reported by Gill et al. (2010), we sought to confirm that our growth factor treatments were in fact eliciting the appropriate cellular responses in HeLa cells. To this end we performed a time course of EGF or PDGF treatment following serum starvation of HeLa cells. Cell extracts were analysed by Western blot using antibodies specific to phosphorylated P44/P42 MAPK as an indicator of receptor tyrosine kinase activation and subsequent downstream signalling (Katz et al., 2007; Murphy and Blenis, 2006; Yoon and Seger, 2006). In serum starved cells treated with either EGF or PDGF, peak detection of phosphorylated P44/P42 occurred 10 minutes after treatment and then decreased over the remainder of the time course (Fig. 3A, 3B). These results are consistent with previously reported time frames for MAPK pathway activation (Gao et al., 2005; Pierce et al., 2001).

**Figure 3.**
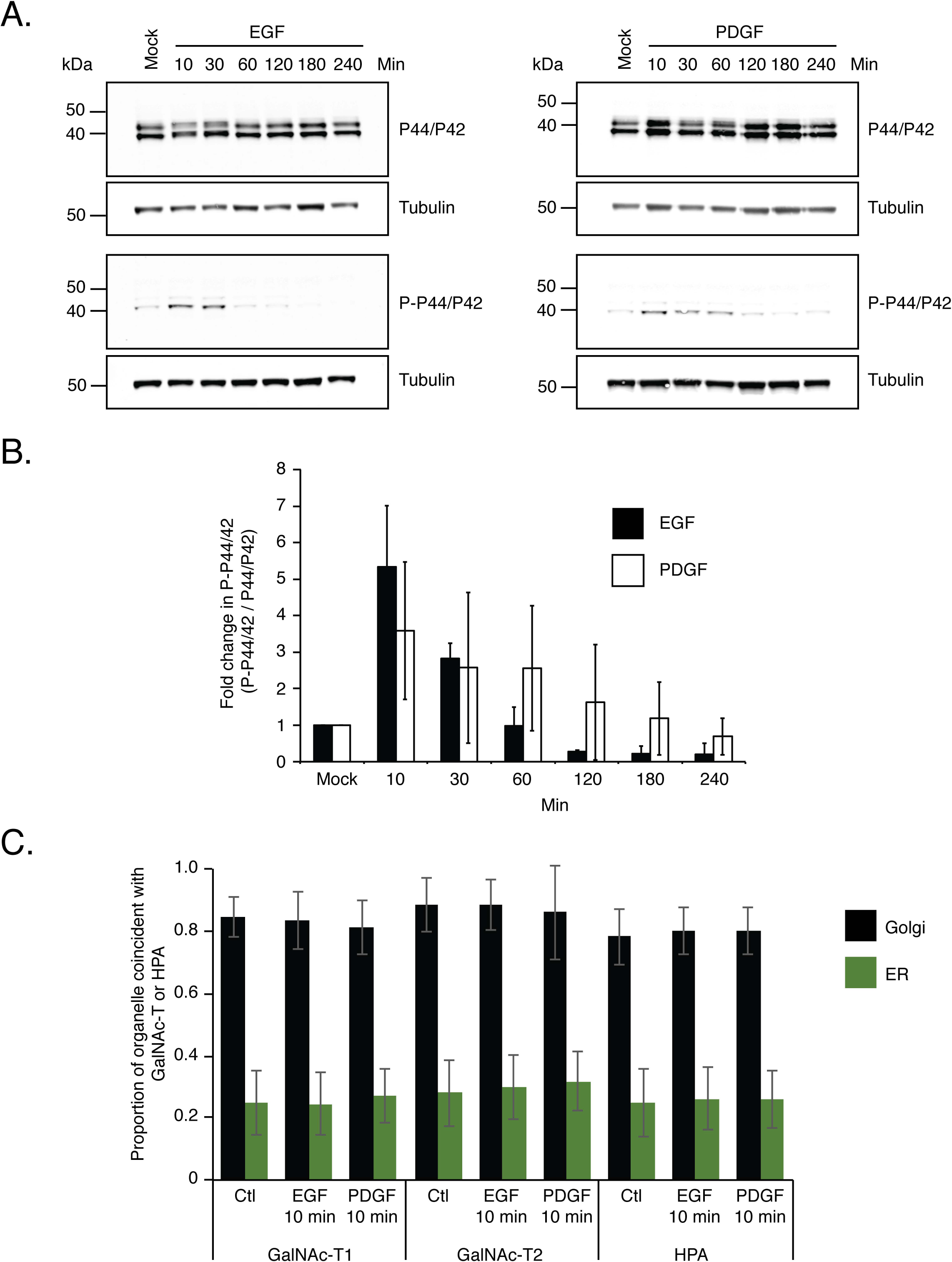
Time course of MAPK activation following growth factor treatments. (A) Serum starved cells were left untreated, treated with 100 ng/ml of EGF or 50 ng/ml of PDGF. At varying time points after treatment cell extracts were prepared and subjected to Western blotting with antibodies specific for P44/P42 or phosphorylated P44/P42 (P-P44/P42) as an indicator of MAPK signalling. The time course revealed that the highest level of P-P44/P42 is detected 10 minutes after treatment. Western blots shown are representative of three independent experiments. (B) Quantification of fold change in detectable P-P44/P42 as a function of time after growth factor treatment. Average fold change in P-P44/P42 is shown with error bars representing the S.D. seen from the quantification of three independent western blots from three separate growth factor treatments. (C) Quantification of amount of GalNAc-T1, -T2 and Tn antigen found in Golgi complex or ER after 10 minutes of growth factor treatments. Manders’ correlation coefficient was calculated on the data set shown in Fig. S2 and Fig. S3. **Values represent the mean ± S.D.**

Our initial growth factor treatments were performed using conditions reported by Gill et al. (2010). Because 4 hour EGF or 3 hour PDGF treatment yielded no change in GalNAc-T Golgi complex localization (Fig 1. and 2) and the fact that there was little detectable MAPK activation at these time points (Fig 3A, 3B), we next determined if GalNAc-T or Tn antigen localization changed after 10 minutes of growth factor treatment, when the highest level of phosphorylated P44/P42 is detected (Fig. 3A, 3B). When serum starved cells were left untreated or treated with either EGF or PDGF for 10 minutes, GalNAc-T1, -T2 (Fig. S2) and Tn antigen (Fig. S3) remained co-localized with TGN46. MCC calculations confirmed that there existed a high degree of coincidence between GalNAc-T1, -T2 and Tn antigen with TGN46 (0.79-0.89 ± 0.1) and low coincidence with calnexin (0.25-0.3 ± 0.1) (Fig. 3C). These data indicate that under conditions when MAPK pathway activation is highest there is still no change in GalNAc-T Golgi complex localization.

## Discussion

Here, we demonstrate that growth factor treatment of serum starved HeLa cells causes no change in the Golgi complex localization of endogenous GalNAc-T1 and -T2, the predominant isoforms expressed in these cells, or Tn antigen, the immediate product of GalNAc-T enzymatic activity. These findings are in direct contrast to what was previously reported in the literature (Gill et al., 2010; Gill et al., 2011). Gill et al. utilized immunofluorescence experiments that simultaneously visualized HPA (a lectin that recognizes Tn antigen)/Tn antibody/giantin (a Golgi complex marker) or HPA/GalNAc-T1/giantin to ascertain if growth factor treatments cause the Tn antigen and GalNAc-T1 to redistribute to the ER. Part of the evidence presented to support redistribution to the ER are images of cytoplasmic regions that are discrete from the areas occupied by the Golgi complex marker. The authors also visualize GalNAc-T1/ERGIC53 (a marker of the ER-to-Golgi complex intermediate compartment)/HPA and HPA/PDI (an ER marker)/giantin to support their hypothesis that GalNAc-Ts are trafficked to the ER after growth factor treatment. Again, insets of cytoplasmic regions are used to support these conclusions. However, Gill et al. did not report simultaneous visualization of a GalNAc-T with a Golgi complex and an ER marker to support their hypothesis. When we performed growth factor treatments using conditions reported by Gill et al. (2010) and simultaneously visualized a Golgi complex marker (TGN46), an ER marker (calnexin) and endogenous GalNAc-Ts or Tn antigen (HPA) no change in GalNAc-T or Tn antigen localization was detected.

To quantify amount of Tn antigen or GalNAc-T1 co-localized with their Golgi complex marker Gill et al. use two different approaches. First, they measured the fluorescent intensity of each marker along a randomly drawn line and looked for similar changes in intensity of each marker along the line. Second, they identified the area of the cell corresponding to the organelle in question, by performing “intensity-based cutoff”, and subsequently determined the degree to which the experimental marker resided within that area. It is unclear if these measurements were made over the entire volume of a cell or only on individual confocal sections. By this approach and the markers used (giantin/HPA and GalNAc-T1/giantin/HPA) the authors would only be able to conclude whether or not Tn antigen and GalNAc-T1 is found in the Golgi complex under the various growth factor treatments. Because simultaneous imaging of an ER marker and GalNAc-T were never performed, it would not be possible to conclude if GalNAc-Ts were located in the ER. Additionally, quantification of experiments involving T1/ERGIC53/HPA and HPA/PDI/giantin were not reported. In the results presented here we directly visualized TGN46, calnexin and either endogenously expressed GalNAc-T1, T2, or HPA. To quantify the degree to which GalNAc-Ts or HPA co-localized with either TGN46 or calnexin we calculated the Manders’ correlation coefficient (MCC) for each combination of markers, visualized over the entire volume of each cell, under the growth factor treatments tested. When comparing two entities, this approach relays the fraction of one channel that co-localizes with the second channel (Dunn et al., 2011; Manders et al., 1993; McDonald and Dunn, 2013). By this measure we confirmed that there was a strong degree of coincidence between both endogenous GalNAc-Ts and Tn antigen with TGN46 and a low degree of coincidence with calnexin under all conditions tested, demonstrating that growth factor treatment did not result in a change in the Golgi complex localization of either GalNAc-Ts or Tn antigen.

Gill et al. (2010) did not report a quantitative measure of EGF or PDGF effect after addition to HeLa cells, except for the immunolocalization of GalNac-T1, HPA or Tn. We therefore cannot rule out the possibility that the effect of EGF and PDGF treatment is temporally different in the cells they employed relative to the HeLa CCL-2, cells obtained from the ATCC. However, to partially address this possibility we also performed our immunofluorescence experiments after 10 minutes of EGF or PDGF treatment, when we detect the highest levels of phosphorylated P44/P42. Our results indicate that regardless of when we measure the localization of GalNAc-Ts they are predominantly found co-localized with TGN46 and thus within the Golgi complex.

We are unable to rule out the possibility that there is an inherent difference between the HeLa cells used by Gill et al. (2010) and those purchased from ATCC. We did initial studies with one sample of cells graciously provided by Dr. F. Bard, and did not find any difference from the results reported here (data not shown). We subsequently learned that a second line of HeLa cells was also used by the Bard lab. At a minimum, our results indicate that the observations termed “GALA”, reported by Gill et al. are not generalizable to all HeLa cells. Thus, care should be taken when exploring the possibility that this reported phenomenon is a global mechanism for regulating mucin-type protein O-glycosylation.

## Materials and methods

### Cell Culture

HeLa CCL-2 cell were purchased from ATCC (American Type Culture Collection (ATCC), Manassas, VA), and maintained in Dulbecco’s Modified Eagle Medium (DMEM) (ThermoFisher Scientific) supplemented with 10% fetal bovine serum (FBS) (ThermoFisher Scientific) and 1% Pen/Strep (Life Technologies, Grand Island, NY) at 37°C in a 5% CO2 incubator.

### Antibodies and fluorescent lectins

For immunofluorescence experiments the following primary antibodies and lectins were used: mouse anti-GalNAc-T1 (un-diluted), mouse anti-GalNAc-T2 (un-diluted)(Mandel et al., 1999); sheep anti-TGN46 (1:500; Biorad); rabbit anti-calnexin (1:200; Abcam); Alexa Fluor^®^ 488 *Helix pomatia agglutinin* (HPA) lectin (5μg/ml; ThermoFisher Scientific). The fluorescently labelled secondary antibodies used were: Cy5 Donkey anti-sheep (1:100; Jackson ImmunoResearch); Alexa Fluor^®^ 488 Donkey anti-mouse (1:1000; ThermoFisher Scientific); Alexa Fluor^®^ 555 Donkey anti-mouse (1:1000; ThermoFisher Scientific); Alexa Fluor^®^ 555 donkey anti-rabbit (1:1000; ThermoFisher Scientific).

### Immunofluorescence

For experiments in which un-transfected HeLa cells were used to quantify changes in the intracellular distribution of endogenous GalNAc-T1, T2 or HPA-labeled species, cells seeded at the desired density onto glass cover slips were grown in DMEM containing 10% FBS. Cells were washed twice with phosphate buffered saline (PBS) and left incubating in serum-free media for at least 16 h. Cells were washed once more and later incubated with or without DMEM containing 0.1% bovine serum albumin (BSA) and human recombinant EGF (100 ng/ml; Sigma Aldrich) or PDGF-bb (50 ng/ml; ThermoFisher Scientific). Cells were washed, fixed with 4% paraformaldehyde (PFA) in PBS for 10 min, washed with PBS, and later incubated in quench solution (NH_4_Q 75 mM, glycine 20 mM) containing 0.1% Tx-100 for 10 min. Cells were blocked in 0.2 % BSA/fish skin gelatin (7mg/ml)/0.05% saponin in PBS for 30 min. Primary mouse antibodies against GalNAc-T1 and T2 were used non-diluted, complemented with 0.5% BSA and with antibodies against calnexin and TGN46 overnight at room temperature. For HPA labelling, cells were incubated with antibodies against calnexin and TGN46 in the blocking solution. Cells were then incubated with the respective secondary antibodies with or without HPA-conjugated to Alexa-488 (5 *μ*g/ml) for 2 h at 37°C. Cells were washed and later mounted with ProLong Gold (ThermoFisher Scientific) reagent and cured overnight before imaging.

### Microscopy and image quantification analyses

Images shown in all figures were obtained with a Nikon A1R+ confocal microscope (Nikon Instruments Inc.) equipped with the following solid state lasers and filters: 488 nm (525/50), 561 nm (600/50) and 640 nm (650LP), using a 60x Plan Apochromat oil objective (NA 1.4) with an image pixel size of 70 nm and a pinhole size of 0.7 airy unit (AU). Image acquisition was performed at room temperature using the NIS-Element imaging software (Nikon Instrument Inc.).

Comparative co-localization analysis of HPA-reactive material or endogenous GalNAc-T1 and GalNAc-T2 with ER marker (calnexin) and Golgi complex marker (TGN46) of control and growth factor-stimulated cells was performed using Imaris (Bitplane, Oxford Instruments, Concord, MA). For each image, a mask is created using the HPA or GalNAc-Ts channel, and then an automatic threshold is applied for each marker channels. The Manders’ coefficient A, corresponding to the proportion of intensity in channel A (ER or Golgi complex) coincident with intensity in channel B (HPA or GalNAc-Ts), is then used to quantify the co-localization of both markers with HPA or GalNAc-Ts.

### Western blot analysis

HeLa cells were plated on 100mm dishes and allowed to recover for ~18 hours. Cells were then washed with PBS and incubated in serum-free DMEM for 16 h. Cells were then treated with DMEM containing 0.1% BSA and human recombinant EGF (100 ng/ml; Sigma Aldrich) or PDGF-BB (50 ng/ml; ThermoFisher Scientific) for 10 min, 30 min, 1 hours, 2 hours, 3 hours or 4 hours. After the indicated times, cells were trypsinized and collected by centrifugation. Following two washed in PBS cells were lysed in 1X Phosphate buffered saline (PBS), 1% Triton X-100 and HALT protease inhibitors (Roche) for 30 min at 4C. Proteins were resolved on 4-12% SDS-PAGE and transferred to nitrocellulose membranes (ThermoFisher Scientific). Membranes were blocked in Block Buffer (Licor) and subsequently incubated with either rabbit anti-P44/P42 MAPK (1:1000; Cell Signaling) or rabbit anti-P-P44/P42 (1:1000; Cell Signaling) and mouse anti-alpha-tubulin (1:5000; Cell Signaling). Following three washed in PBS +0.1% Triton-X100 membranes were incubated with IRDye^®^ 680 donkey anti-mouse (1:5000; Licor) and IRDye^®^ 800 donkey anti-rabbit (1:5000; Licor). Membranes were subsequently washed with PBS +0.1% Triton-X100 followed by PBS immediately prior to data acquisition with a Licor Odyssey scanner. Quantification of Western blot band intensity was performed using ImageJ. Fold change in P-P44/P42 was calculated by first normalizing band intensity for either P44/P42 or P-P44/P42 against the alpha tubulin signal from the mock lane in each blot. Next, the density in the normalized mock lane was divided by the normalized density of each time point to yield a fold change. Lastly, the fold change in P-P44/P42 was divided by the fold change in P44/42 for each respective time point. The average values for each time point were calculated from three independent western blots from three separate growth factor treatments and were graphed using Microsoft Excel. Error bars represent standard deviation.

### RT-qPCR

Total RNA were extracted from HeLa ATCC cells using the RNeasy micro kit and the QIAshredder spin column (Qiagen, Valencia, CA) for homogenization, following manufacturer protocol. Then 1mg of RNA was used to produce cDNA using iScript™ Reverse Transcription Supermix (Biorad, Hercules, CA). RT-qPCR was performed using Taqman^®^ Gene expression assay (ThermoFisher Scientific, Waltham, MA) following manufacturer procedure. The PCR was performed using TaqMan^®^ Universal PCR Master Mix (ThermoFisher Scientific, Waltham, MA), with 50 ng of cDNA and the following probes: Actin B (Hs99999903_m1) was used as a positive control, and used to normalization for the relative expression of GalNAc-T1 (Hs00234919_m 1), GalNAc-T2 (Hs00189537_m1), GalNAc-T3 (Hs00237084_m1), GalNAc-T4 (Hs00559726_s1), GalNAc-T5 (Hs00294826_m1), GalNAc-T6 (Hs00200529_m1), GalNAc-T7 (Hs00213624_m1), GalNAc-T8 (Hs00213610_m1), GalNAc-T9 (Hs00222357_m1), GalNAc-T10 (Hs00213903_m 1), GalNAc-T11 (Hs00757865_m1), GalNAc-T12 (Hs00226436_m1) and CD3e (Hs01062241_m1).

## Acknowledgments

We thank Kanishka Patel for help with experiments, Dr. George Patterson for advice on quantification of confocal microscopy data and providing reagents, and Dr. Kenneth Yamada for critical reading of earlier versions of the manuscript. We thank Dr. Ulla Mendel for providing antibodies to endogenous GalNAc-T1 and -T2, and for her review of the manuscript. We thank Drs. Kelly Ten Hagen and George Patterson for reviewing multiple versions of the manuscript. We thank Dr. Frederic Bard for providing us with additional details of his work and a sample of HeLa cells from his lab. This work was supported by the Intramural Program of NIDCR, NIH.

## Abbreviations List

GalNAc-T: UDP-GalNAc polypeptide:N acetylgalactosaminyltransferase
GalNAc: N-acetylgalactosamine
ER: Endoplasmic reticulum
EGF: Epidermal growth factor
PDGF: Platelet-derived growth factor
MAPK: Mitogen-activated protein kinase
NA: Numerical aperature
AU: Airy unit
MCC: Manders’ correlation coefficient

